# Longitudinal dynamics of microvascular recovery after acquired cortical injury

**DOI:** 10.1101/2022.03.09.483613

**Authors:** Xiaoxiao Lin, Lujia Chen, Amandine Jullienne, Hai Zhang, Arjang Salehi, Mary Hamer, Todd Holmes, Andre Obenaus, Xiangmin Xu

## Abstract

Acquired brain injuries due to trauma damage cortical vasculature, which in turn impairs blood flow to injured tissues. There are reports of vascular morphological recovery following traumatic brain injury (TBI), but the remodeling process has not been examined longitudinally in detail after injury *in vivo*. Understanding the dynamic processes that influence recovery is thus critically important. We evaluated the longitudinal and dynamic microvascular recovery and remodeling up to 2 months post injury using live brain miniscope and 2-photon microscopic imaging. The new imaging approaches captured dynamic morphological and functional recovery processes at high spatial and temporal resolution *in vivo*. Vessel painting documented the initial loss and subsequent temporal morphological vascular recovery at the injury site. Miniscopes were used to longitudinally image the temporal dynamics of vascular repair *in vivo* after brain injury in individual mice across each cohort. We observe near-immediate nascent growth of new vessels in and adjacent to the injury site that peaks between 14-21 days post injury. 2-photon microscopy confirms new vascular growth and further demonstrates differences between cortical layers after cortical injury: large vessels persist in deeper cortical layers (>200 μm), while superficial layers exhibit a dense plexus of fine (and often non-perfused) vessels displaying regrowth. Functionally, blood flow increases mirror increasing vascular density. Filopodia development and endothelial sprouting is measurable within 3 days post injury that rapidly transforms regions devoid of vessels to dense vascular plexus in which new vessels become increasingly perfused. Within 7 days post injury, blood flow is observed in these nascent vessels. Behavioral analysis reveals improved vascular modulation after 9 days post injury, consistent with vascular regrowth. We conclude that morphological recovery events are closely linked to functional recovery of blood flow to the compromised tissues, which subsequently leads to improved behavioral outcomes.

## Introduction

The vascular system is comprised of an organized network of arteries, capillaries and veins which are required to efficiently deliver oxygen and nutrients and remove waste products from metabolically active brain tissues^1^. Small vessels/microvessels including arterioles, venules and capillaries (<100μm in internal diameters) are essential to the function of every cell in the central nervous system. Brain microvessels are primarily comprised of endothelial cells along with pericytes and astrocytes, and other cell types. Larger vessels are also ensheathed in smooth muscle cells to aid in contractility^2^. Microvascular restoration processes are critical for post-brain injury tissue preservation and neural circuit reorganization^3, 4^. Due to technical difficulties in measuring small blood vessels in the living brain, scant research has been directed to understanding how cortical vascular networks respond dynamically *in vivo* over time after cortical brain injury, such as the events that occur following traumatic brain injury (TBI)^5, 6^.

Earlier studies have reported apparent structural recovery of brain vasculature following TBI^3, 5, 7^. But these data are restricted to temporal and regional snapshots that collectively are the basis for inferring the progression of post-injury events rather than providing a dynamic highly resolved continuum of events in a single brain region following a single injury. There are no definitive longitudinal dynamic spatio-temporal studies of how the cerebral vasculature, and specifically small vessels, respond to TBI^4, 8^. Detailed angio-architecture and functional recovery of the vascular system following brain injury has been poorly studied, and earlier research has focused on the acute injury period rather than longer term recovery processes^4, 8–10,11, 12^. Thus, our understanding of how brains respond to injury and how we monitor the effectiveness of new treatment strategies is limited.

To address this gap, new studies using high-resolution longitudinal imaging techniques are critical for obtaining a better understanding of microvascular restoration mechanisms. We have applied lightweight, head-mounted, miniaturized microscopes (“miniscopes”)^13–15^ and *in vivo* two-photon (2P) imaging with anatomical vessel painting of brain vasculature to examine dynamic microvascular restoration following TBI and aspiration injury in the cerebral cortex of mice. Two distinct models of brain injury were employed to demonstrate similar restorative mechanisms, albeit with different timelines. TBI models have been used to model moderate and severe penetrating injuries^4, 8, 16^ whereas cortical aspiration brain injury models are used typically to assess neural and vascular repair^17, 18^. Miniscope imaging allows for repeated high resolution and high-speed longitudinal imaging of the vasculature and associated cells, as well as determination of blood flow in the same mouse following cortical injury. There are considerable interpretive and technical advantages for longitudinal studies within animals that permit dynamic measures of post injury responses, coupling microvascular restoration with behavioral assessments.

We set out to test the hypothesis that focal cortical injury which leads to dramatic reduction in vascular structure and function, initiates *de novo* vascular growth and vessel pruning to form new semi-functional vascular networks. We determined the time course and spatial patterns of vascular re-growth and blood flow dynamics after focal cortical injury. We report on these foundational processes of vascular restoration following cortical brain injury.

## Materials and methods

### Animals

All experiments were conducted in accordance with the National Institutes of Health guidelines for animal care. All animal use was approved by the Institutional Animal Care and Use Committee and the Institutional Biosafety Committee of the University of California, Irvine. Mice had access to food and water in their home cages with lights maintained on a 12 h light/dark cycle (lights on at 6:30 am, off at 6:30 pm). Male Tie2-Cre; Ai9 mice (*n*=18, 10 for miniscope, 8 for 2P; 3-4-month-old) expressing tdTomato (red fluorescent protein) in vascular endothelial cells were obtained by crossing Tie2-Cre transgenic mice with Ai9 mice and used for miniscope and 2P recordings. The Tie2-Cre transgene has the mouse endothelial-specific receptor tyrosine kinase (Tek or Tie2) promoter directing expression of Cre recombinase in vascular endothelial cells (JAX #008863). The Ai9 mouse is a Cre reporter mouse strain on the C57BL/6J genetic background (JAX #007909), and Ai9 mice express robust tdTomato fluorescence following Cre-mediated recombination. Vessel painting experiments were performed in male C57BL/6J (*n*=6) and in Tie2-GFP mice (*n*=4) (8-12-week-old, Jackson laboratories). Experimental time points were selected based on previously published data from our laboratories and available literature^3, 5, 19^. A separate cohort of Tie2-Cre; Ai9 mice (*n*=13, 7 males and 6 females, 3-4-month-old) was utilized for behavioral experiments in relation to outcomes of the functional restoration of the vasculature.

### Brain Injury Models: Aspiration and traumatic brain injury

The brain injury models comprised two groups: cortical aspiration and cortical contusion injury (CCI). CCI was induced in mice as previously described^20^. Briefly, mice were anesthetized with isoflurane (3% induction, 1-2% maintenance, VetEquip Inc., Pleasanton, CA, USA) and placed on a heating pad to maintain body temperature (37°C). Carprofen (5 mg/kg) was injected subcutaneously before surgery for pain relief and to reduce inflammation. The mouse head was shaved and cleaned with 70% ethanol and swabbed with povidone-iodine. A midline incision was performed, and skin and connective tissue were pushed aside to access the skull. A burr drill was used to roughen the surface of skull to facilitate application of dental cement for miniscope experiments (see below). After brain injury induction, the surgical wound was sutured closed and taped with tissue adhesive (3M Vetbond, St. Paul, MN).

#### Traumatic brain injury

A 2.3mm-diameter trephine was used and access to the sensorimotor cortex was positioned 1.25mm posterior from Bregma and 1.25mm lateral from the sagittal midline. A CCI was induced by an electromagnetic impactor (Leica Microsystems Company, Richmond, IL) with a flat stainless-steel tip (speed: 5 m/sec, diameter: 1.5mm, depth: 1mm; impact diameter for vessel painting mice: 5mm, impact diameter for miniscope/2P mice: 3mm) was used to induce a cortical contusion. Excess blood was carefully swabbed from the injury site. Minimal herniation was observed in mice after CCI in vessel painted, miniscope and 2P experimental mice. In our TBI experiments we did not note any overt tissue loss during the subacute period; implantation of the GRIN lens could possibly mask minimal tissue loss.

#### Aspiration injury

A 2mm-diameter hole was drilled above motor cortex (AP: 1.54mm; ML: 1.8mm; DV: 0.2-0.5mm from Bregma) and the underlying cortex was lightly aspirated (~2mm diameter and 500μm depth). Cortical tissues were rinsed with saline until bleeding stopped.

#### Controls/Shams

Appropriate shams and controls were utilized for each type of experiment. Controls and shams underwent the identical procedures with the absence of brain injury.

### *In vivo* miniscope imaging

#### GRIN lens implantation

Miniscope gradient-index lens (GRIN) implantation surgery has been previously described^14^. Mice were implanted with lenses immediately after CCI or aspiration brain injury for *in vivo* vascular imaging. The lens holder and GRIN lens (0.25 pitch, 0.55 numerical aperture, 1.8-mm diameter and 4.31 mm in length) were attached to a stereotaxic apparatus. GRIN lens was then gently lowered into the cortex (~200-300μm) with Krazy glue applied around the lens sealing the craniotomy and exposed tissue. The microendoscope was secured to the skull with dental cement (Lang Dental Manufacturing, Wheeling, IL, Item: 1304CLR) to stabilize the miniscope. After dental cement dried, a thick layer of Kwik-Sil (Word Precision Instruments, Sarasota, FL) was applied on the top of the lens to protect it from physical damage. After implantation, the mouse recovered in a clean cage before being returned to its home cage.

#### Miniscope imaging

Please refer to our previous publications^14, 21^ and www.miniscope.org for technical details of our custom-constructed miniscopes. Vascular regrowth was recorded daily using miniscopes with high spatial resolution (~0.9μm per pixel, field of view: 700μm x 450μm at 60Hz) under three conditions: anesthesia, awake and running states. FITC-conjugated dextran (Item: FD2000S; 2000kDa Sigma-Aldrich, St. Louis, MO) was dissolved in purified water at a concentration of 50 mg/ml and protected from light during the preparation. The mouse was anesthetized with isoflurane (3% induction, 1-2% maintenance) and received 150-200μl FITC-conjugated dextran (10mg/ml via tail vein injection. Blood flow was recorded for a 2 min duration.

### *In vivo* 2-photon (2P) microscopic imaging

#### Chronic cranial window surgery

To perform 2P microscopic imaging of vascular regrowth, mice underwent a cranial window implantation surgery. The mouse was anesthetized with isoflurane (3% induction, 1-2% maintenance) and body temperature was controlled by a thermoregulated heating pad. Artificial tears ointment was applied to protect the eyes. Carprofen was injected subcutaneously (5 mg/kg). Access to the skull was as described above. After the exposed skull was completely dry, a thin layer of Krazy glue was applied and allowed to harden. A 2.8mm diameter cranial window mark was drawn (AP: 1.54mm; ML: 1.8mm from Bregma). Acrylic was applied to the exposed skull and on a U-shaped head fixation bar, centered at the bottom of the demarcated cranial window location (~3-4 mm). A 2.8mm-diameter trephine opened a cranial window, followed by a CCI (described above). Once bleeding stopped, a 4mm-diameter glass coverslip was affixed to the skull covering the craniotomy. A thin layer of dental cement was used to glue the cover glass and head bar together.

#### Longitudinal vascular imaging

Tie2-Cre; Ai9 mice in the brain injury group were imaged using a Neurolabware 2P imaging system with Coherent Chameleon Discovery laser to assess vascular function and vessel regrowth. Mice underwent 2P imaging daily and the control group was imaged twice a week. Animals were lightly anesthetized with isoflurane (3% induction, 1-2% maintenance) during imaging and body temperature was maintained using a heating pad. A bright field image was captured with a 4X objective to assess the overall vascular regrowth. Then, mice were imaged with the 2P system to capture the tdTomato-expressing vascular structure using a water immersion 16X objective with 1000 nm 2P laser excitation wavelength (line/frame 512; frame rate 15.49 Hz). Z-stack images or Optotune scans (frame rate 20, period 30) were acquired using Scanbox acquisition software (Neurolabware). FITC-dextran (50mg/ml) was injected in tail vein or via retro-orbital injection (30G needle) after baseline recording. A Z-series of blood vessel and flow images (2 or 5μm per step, 30 steps) at a depth of 100–300 μm from the pial surface were captured using both green (FITC, 920nm 2P laser excitation) and red channels.

Different magnifications (1X, 1.7X, 3.4X, 5.7X, 8X) were used to image the individual blood vessels for filopodial growth and to acquire a video capture of fluorescent labeled blood flow. For longitudinal imaging, the same imaging fields and zoomed-in individual vessels at different time points after initial baseline imaging were identified based on the anatomical location and patterns of the vessels. The depth and orientation of the vessels were carefully adjusted so that the identical vessels were imaged, using the same imaging parameters, including pixel size and scan speed. The animals were allowed to recover after each imaging session on a heated pad and then returned to their home cages when ambulatory. Filopodia growth was assessed temporally in a new set of mice (n=4) for in-depth temporal analyses using 2P microscopy. Mice were imaged every 2d after TBI induction for the first 14d and then at 21 and 28dpi. While apparent filopodia appeared *in vivo* at <5d post injury, they became less apparent as vessel density increased. Our analyses included quantification of the density of vessel components across imaging time points and different cortical depths. Briefly, background noise was removed using a region that did not contain vessel segments and density was then measured from the entire image using ImageJ. Data from each mouse were plotted for rate of growth and then normalized as a percent growth using the first imaging session as control.

### Blood flow velocity analysis

To quantify the blood flow velocity, we investigated the blood flow profile along the vessel central line. We split the central line into discontinuous segments, each segment being 60 pixels long (1 pixel=1μm). For each segment we created a line scan profile of blood flow. Red blood cells (RBC) did not have fluorescence and thus appeared dark. Movement of RBCs created dark streaks in the line scan profile that represented blood flow over time. The speed of blood flow can be achieved from the slope of these streaks (distance/time)^22^. First, the line scan profile was smoothed with a 3X3 (pixel) gaussian kernel and then binarized using a threshold equal to 0.5X maximum intensity. In the binarized line scan profile, the pixels forming bright patterns were identified and the slope of the pattern was estimated from the regression line of the pixels. Slopes with a value equal to 0 or infinity were discarded. Due to noise contamination, detected slopes may have both positive and negative values. We chose the sign that most of the slopes possessed with the other signs discarded. The remaining slope values were averaged to achieve the flow speed of the vessel segment. The flow speed of the central line was used to represent the flow speed of all neighborhood vessel areas. To create the flow speed colormap, gaussian smoothing was applied to the original flow speed map to smooth the speed transition between segments. The distribution of blood flow speed was calculated from all vessel segments in the original flow speed map (unsmoothed) and was fitted to a Gamma distribution.

### Vessel painting and analyses

The vessel painting technique is based on the ability of the fluorescent dye 1,1’- dioctadecyl-3,3,3’3’-tetramethylindocarbocyanine perchlorate (DiI; Life Technologies, Carlsbad, CA) to bind to lipid membranes. We have described this methodology in detail^23^. Briefly, mice were anesthetized using an intraperitoneal injection of ketamine/xylazine (90/10mg/kg) and several minutes later the chest cavity was opened to expose the heart. Vessel painting was performed by injecting a solution of DiI (0.3mg/mL in phosphate buffered saline (PBS) containing 4% dextrose, total volume 500μL) into the left ventricle of the heart, followed by a 10mL PBS flush and finally a 20mL 4% paraformaldehyde (PFA) perfusion. Solutions were delivered using a peristaltic pump (8.4 mL/min). Brains were then extracted and post-fixed in 4% PFA for 24 hr and were then rinsed and stored in PBS at 4°C until imaging. Brains were imaged using a fluorescence microscope (Keyence BZ-X700; Keyence Corp., Osaka, Japan). Images of the entire axial brain surface were acquired using the 2X magnification and 1mm depth of field (25.2mm pitch, 40 slices). Resultant images were reconstructed using the XY-stitching and Z-stack features of Keyence software (BZ-X800 Analyzer, Version 1.1.1.8), including haze reduction. Analysis for classical vessel features was performed on entire axial brain images using AngioTool software (Version 0.6)^24^. Miniscope and 2P images of vessels were also analyzed using identical software setting. Quantitative vessel features included vessel density, number of junctions and vessel length.

### Behavioral testing

All brain-injured and control/sham mice underwent habituation handling in the behavioral testing room prior to actual testing. Baseline testing was performed for both foot-fault and balance beam tests in mice one week prior to testing after brain injury. Behavioral tests were acquired at discrete days post-injury in injured and control mice (2, 7, 14, 20, 27dpi).

#### Foot-fault

Foot-fault testing was performed on an elevated wire mesh rack, measuring 14×41×21cm (height/length/width) with 21 rungs evenly spaced apart (1.6cm). Each animal was placed in the center of the rack and its movement was video recorded for a period of 3min. A foot-fault was recorded when a mouse paw slipped through the mesh. Scoring was performed manually by tracking foot-faults for each paw individually. A foot-fault score for each animal was calculated based on the total number of foot-faults/paw over the distance traveled.

#### Balance beam

Balance beam testing was performed on an elevated (61 cm high) acrylic beam balance (61cm long) with a 0.65cm walkway for animals to traverse. Each animal was placed at the midpoint of the beam and allowed to walk unrestricted in either direction for 3min while being video recorded. The number of slips for each paw, total time moving, and distance traveled were manually scored. Balance beam scores for each mouse was calculated as a ratio of total slips/paw over total distance traveled.

### Statistical analysis and reproducibility

Researchers were blinded to experimental conditions for all data quantification. Data were analyzed as described above and presented as mean ± SEM. All data were graphed in Graphpad Prism 8.0. Statistical tests utilized are detailed within the text and in the figure legends. Briefly, temporal responses were analyzed using a one- or two-way ANOVA followed by Tukey post-hoc testing. Group responses were modeled using linear regressions and plotted as mean with 95% confidence intervals (CI). In the cortical aspiration data, linear regressions in the control mice were extended from 30dpi to 60dpi to better model the temporal response (see Fig. 3). Repeated measures statistical testing was performed in within-subject acquired data. A *p*-value of 0.05 was considered statistically significant. All experiments were performed with a three or more experiments or biological replicates, unless otherwise noted.

### Data availability

All data that were generated or analyzed during this study are provided in this publication, including supplementary information files. Raw data are available upon reasonable request from the authors.

## Results

### Vascular loss followed by de novo microvessels regrowth after cortical injury

We used a novel DiI vessel painting approach to label vascular endothelium after cortical brain injury^23^ to anatomically map the temporal progression (1, 7, 14 and 30dpi); n=5-6 mice/ time point) of vascular recovery after a single impact of brain injury (Fig. 1). Compared to sham controls, a focal impact injury to sensorimotor cortex evoked a significant loss of cortical vasculature at superficial cortical layers at the injury site at 1dpi (Fig. 1A, B). The vessel void was characterized by discontinuous vessels, vessel fragmentation and in some severe cases, extravasation of the DiI label into the surrounding parenchyma (Fig. 1B, C). Over the course of the next 30dpi, vessel density dynamically increases with progressive significant increases in vessel number (7dpi), peaking at 14dpi, then returning to control levels by 30dpi (Fig. 1A-C). Post-injury vessels at 30dpi are morphologically distinct from pre-injury control vessels: they are more tortuous and show non-uniform vascular density (Fig. 1C; see also Supplementary Fig. 1).

**Fig. 1.**
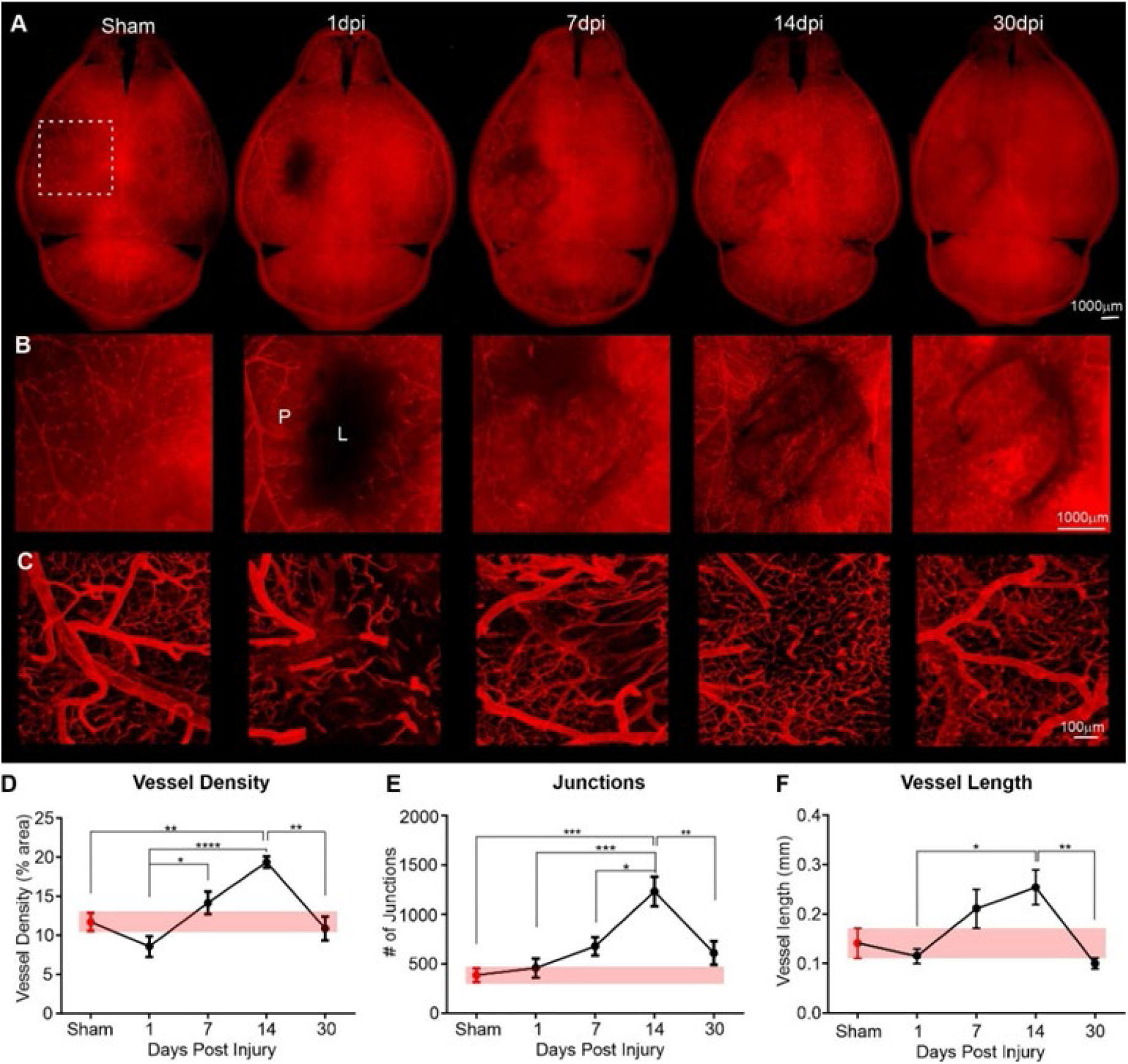
Sequence of vascular morphological recovery following focal contusion brain injury in mouse cortex. A) Wide field epifluorescent images of the whole brain (A), expanded view of the injury site (B), and confocal microscopy in the perilesional area (C). At 1-day post injury (dpi) there is a dramatic loss of vessels at the lesion (L) site. The lesion area is refilled with small blood vessels and other cells over the 30d time course. Revascularization appears to be most prominent at 7-14dpi. Confocal images (C) reveal that new vessels exhibit distinct morphologies as compared to sham/naïve vessels, with abnormally increased branching and smaller vessel structures. D) Overall vascular density in the perilesional region (P) of the injured hemisphere is reduced at 1dpi but peaks at 14dpi, then returns to sham levels (red bar) by 30dpi. E) Lesion induced changes in vascular junctions follow a similar time course to vessel density. F) Post-lesion average vessel length also slowly increases, reaching a maximal length at 14dpi, followed by a return to sham (red bar) levels. (n=5-6/time point; * p<0.05, ** p<0.01, *** p<0.001, **** p<0.0001, one-way ANOVA with Tukey’s multiple comparisons post-hoc tests)

To confirm qualitative visual assessments, we quantitatively measured vascular features, including vessel length, density, and numbers of junctions. Superficial layer vessel density in the peri-injured site declines at 1dpi but then significantly increases over the next 14d with a return to control vascular density levels by 30dpi (Fig. 1D). Increased vessel density is accompanied by a highly significant ~3-fold increase in the number of vessel junctions, consistent with vessel proliferation (Fig. 1E). Vessel length progressively increases by ~2 fold at 14dpi, then decreases to control values at 30dpi, suggesting vascular pruning or regression processes at this later stage (Fig. 1F). Thus, vessel painting documents the temporal loss and subsequent restoration of cerebral vasculature following a single impact of brain injury.

### In vivo longitudinal imaging of post-injury microvascular recovery

The temporal changes in angioarchitecture visualized with vessel painting are static snapshots from different animals. To visualize post-injury vascular dynamics at greater temporal resolution in the same animals, we applied *in vivo* miniscope imaging (Fig. 2A-E) with a GRIN lens implanted into the injury site in two types of cortical injury: 1) TBI using a CCI model (Fig. 2), and 2) focal aspiration of superficial cortex (Fig. 3). The implanted GRIN lens and fixed miniscope baseplate allow reliable, repeated imaging of the same group of vessels over extended periods of time. The head-mounted miniscope was magnetically attached to the baseplate without a permanent fix. Under our experimental conditions, the miniscope has a 700μm x 450μm field of view with a resolution of 752 pixels x 480 pixels (~0.9μm per pixel) and measures an approximate depth of 50-100μm down from the GRIN lens surface^14^. The lens was immediately implanted after injury and miniscopes were used to longitudinally image *in vivo* events continuously up to two months or longer (Figs. 2, 3). In control/sham mice the GRIN lens was fixed to the cortical surface without damage to the superficial vascular network. In control mice, a dense vascular network is observed on the cortical surface that is composed of large vessels, as well as small microvessels (Fig. 2D). Quantification of the control/sham vasculature reveals an average vessel density in terms of cortical coverage (32.99 ± 1.12% of the imaging field of view, n=4), average total number of junctions of 120.50 ± 14.57 and average vessel length of 214 ± 23.33 mm averaged across all time points (1, 4, 7, 13, 21 and 30dpi). There are no significant differences in any control mice vascular parameters (p=0.34-0.62). Cortical injury in mice results in a similar pattern of vessel re-growth as observed with vessel painting (compare Figs. 1, 2E, 3A). Importantly, repeated longitudinal miniscope imaging documents vascular regrowth into functional vascular networks when labeled by circulating fluorescein-labeled dextran (Fig. 2E, 3A; Supplementary videos 1 and 2).

**Fig. 2.**
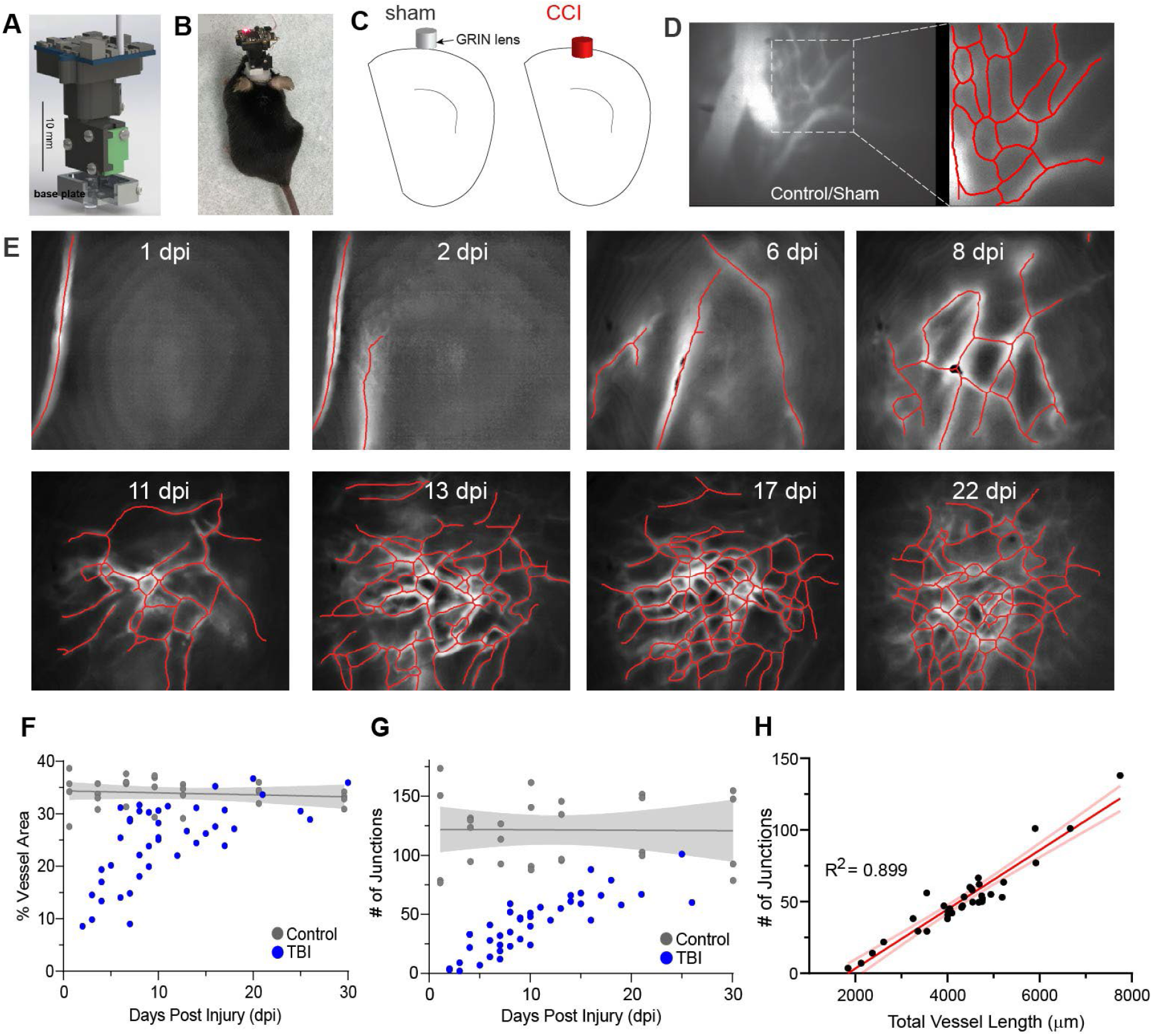
Longitudinal *in vivo* miniscope imaging of temporal and spatial patterns of cortical microvascular networks following cortical contusion injury. A) Schematic of our miniature microscope for microcirculation imaging (details at http://miniscope.org/). B) A mouse fitted with the head mounted microscope. C) Schematic locations of GRIN lens implantation in sham and cortical contusion injury (CCI=TBI) mice. D) Miniscope imaging of the cortex of control/sham mice reveals a diversity of vessel sizes, ranging from large cortical vessels on the surface of the cortex along with dense plexus of smaller vessels. Microvascular networks were visualized by intravenous injection of fluorescent-labeled dextran. E) Longitudinal tracking of cortical angioarchitecture from the same mouse spanning 1 to 22dpi in the same field of view illustrates the progressive repair of the vasculature, following TBI. Vessel center lines delineated by software assist visualization (red lines). F) Vessel density over the 31dpi observation rapidly increases over the first 14d and then plateaus in the TBI model (n=4) with control/sham (n=4) mice having a stable vessel coverage (gray line with 95%CI). G) The number of junctions also increases with time in all TBI mice (n=4). Control/sham mice have a stable number of vascular junctions (gray line, 95% CI; n=4).H) A strong linear correlation (R^2^=0.899, red line) is observed where the number of junctions increases linearly with the increasing total vessel length (n=4). The 95% confidence intervals are also plotted.

**Fig. 3.**
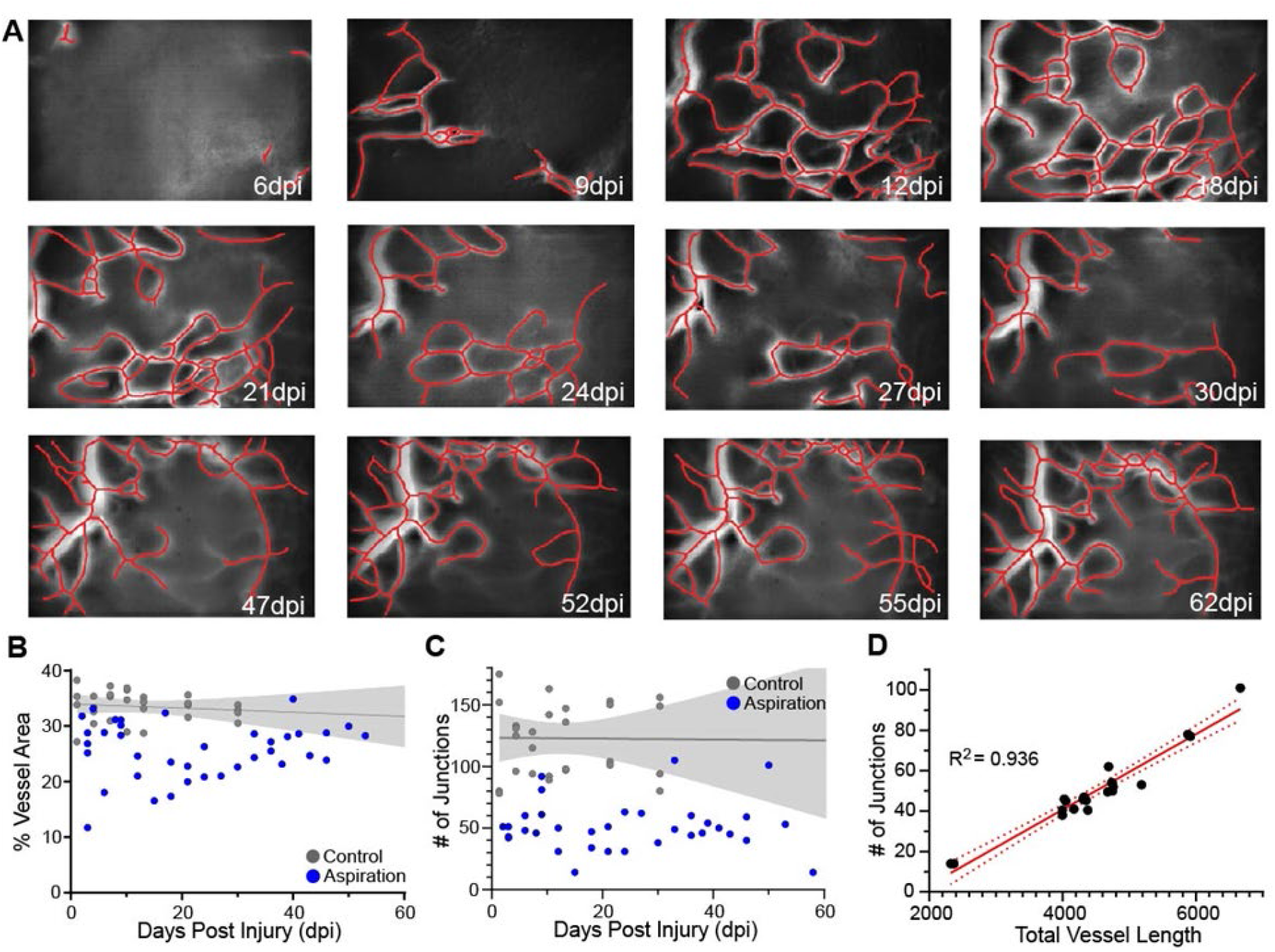
Microvascular regrowth in the aspiration cortical injury model does not follow the exact pattern as observed in the TBI model. A) Longitudinal tracking of cortical micro-vessels from the same mouse spanning 6 to 62dpi acquired from the identical field of view illustrates the progressive increase in vessel density after focal aspiration injury. Vessel center lines delineated by software, assist visualization (red lines). B) The overall vessel density over the time course up to ~60dpi in the aspiration injury model (n=4). Control/sham (n=4) mice have a stable vessel coverage up to ~30dpi (gray line = linear regression with 95%CI fitted to 60dpi). C) The pattern of the number of junctions approximately mimics that of the vessel density in (B). Control/sham mice have a stable number of vascular junctions over 30dpi (gray line = linear regression with 95%CI fitted to 60dpi n=4;). D) Similar to the TBI model, there is a strong linear correlation (R^2^=0.936, red line) where the number of junctions increases linearly with the increasing total vessel length (n=4). The 95% confidence intervals are also plotted (red dashed line).

Vascular regrowth in TBI (n=5; Fig. 2) and aspiration cortical injury (n=5; Fig. 3) models do not follow identical patterns. Vessel density from repeated measurements in a cohort of TBI mice show rapid increases in vessel density peaking at ~20dpi (Fig. 2F), where each measurement for all mice is plotted. In TBI animals this vascular growth appears to plateau between 10-30 days (Fig. 2E). The number of vascular junctions also increases progressively immediately after injury, but with a more delayed and gradual onset (Fig. 2G). There is a strong correlation between total vessel length and number of vessel junctions (R^2^=0.899) in TBI mice, further pointing to a synergistic generation of a renewed and potentially functional vasculature. One apparent difference between post-injury responses as measured by static analysis (Fig. 1) versus longitudinal imaging (Fig. 2) is that we do not see clear evidence for vascular regression for the parameters of vessel area and number of junctions between 14 and 30dpi when measured by miniscope imaging (Fig. 2F-H). The most parsimonious explanation for this difference is that with miniscope longitudinal imaging, we can precisely follow the entire history of each post-injury vessel within each mouse over time within the lesion site, while for static measurements based on vessel painting, we can only infer from each “snapshot” of individual animals at the perilesional site.

Aspiration cortical injury, as noted above has a different time course for vascular recovery (Fig. 3), but there is a similar vascular recovery (Fig. 3A). Initially, there is not an overt loss of vessels (some variability between mice exists) but by 14dpi there is an overall decrease in vascular density that slowly recovers over the next 58dpi (Fig. 3B). The temporal vessel junction evolution is more variable in this model (Fig. 3C). Like the TBI cohort, we observe a strong correlation in the number of junctions and total vessel length after aspiration injury (R^2^=0.936) (Fig. 3D). While temporal differences exist between the two models, both types of cortical injury demonstrate vascular regrowth and functional recovery (see below).

### TBI results in increased vascular remodeling in superficial cortical layers

To identify changes in vascular features at different cortical depths, we used 2-photon (2P) microscopic imaging (Fig. 4A). *In vivo* 2P imaging of cortical vasculature labeled by fluorescein-labeled dextran showed that the control cortex exhibits a vascular network comprised of a mixture of large and small vessels (Fig. 4B), similar that seen in miniscope imaging data (Fig. 2D). 2P imaging reveals a remarkably altered topology of cortical vasculature in TBI mice, even at 25dpi (Fig. 4C). Further examination of the post-injury vascular network at different depths reveals striking differences in the remodeled vascular network (Fig. 4D; 25dpi). Post-injury vascular density is reduced, but changes in the heterogeneity of post-injury vascular diameters are apparent (Fig. 4C) as compared to control (Fig. 4B). The upper layers of post-injury cortex exhibit a higher density of small tortuous vessels with vessel diameters ranging from 4.8 to 19.4μm (Fig. 4D, upper cortex). At deeper layers into the cortical parenchyma, there is a preponderance of large vessels (Fig. 4D, lower cortex) whose vessel diameters range from 11.8 to 88.8μm with 54% of the vessels having a diameter >40μm (see Supplemental Fig. 1E). This suggests more extensive post-injury vascular remodeling in superficial cortical layers that results in re-growth of a plexus of smaller vessels that occurs around 3 weeks after injury.

**Fig. 4.**
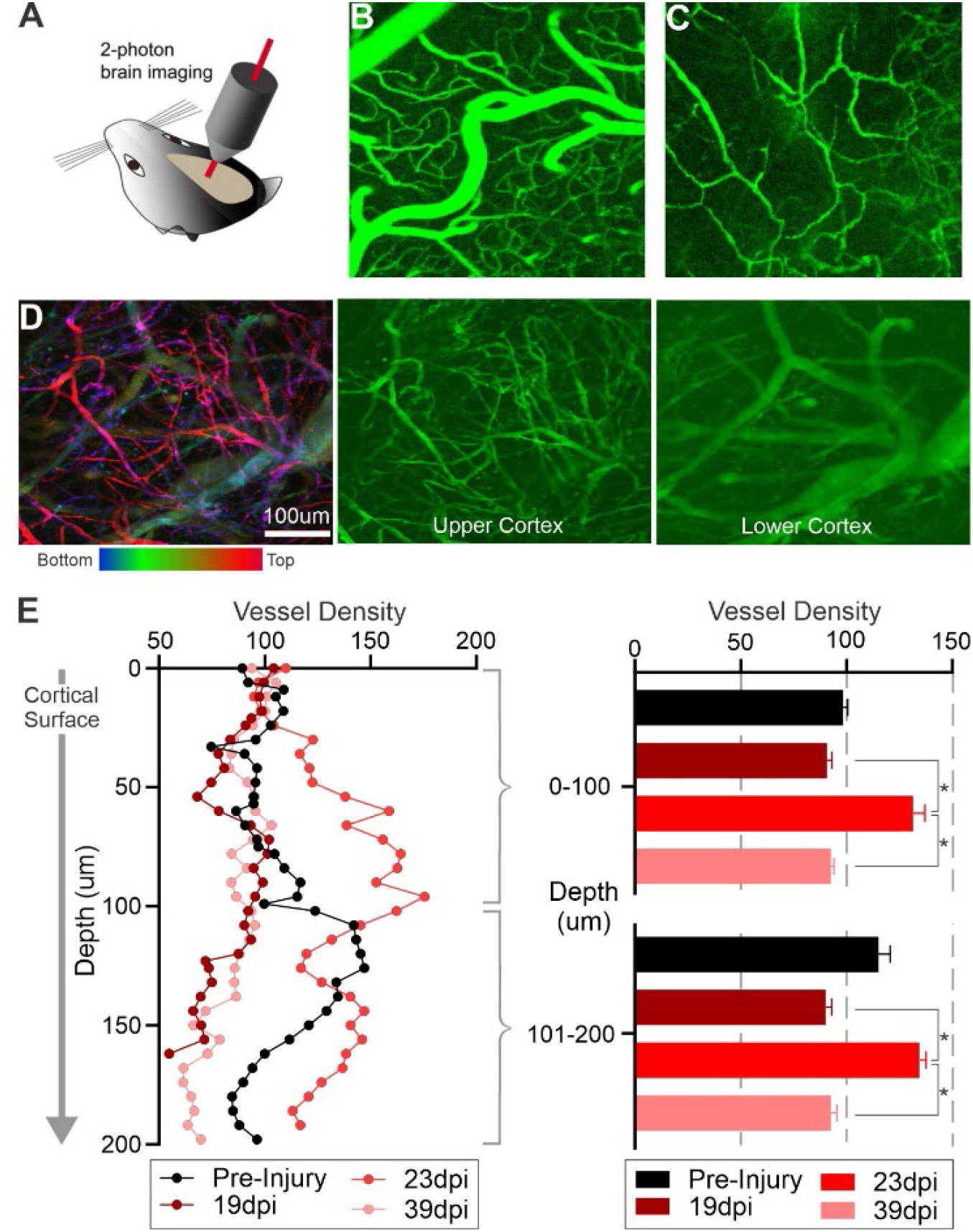
*In vivo* two-photon imaging shows differential microvascular remodeling at different cortical depths in the TBI model. A) Schematic of 2P structural imaging of blood vessels in the mouse brain *in vivo* through thin skull preparation. B) 2P laser scanning microscopy in conjunction with labeling of the blood plasma with Fluorescein isothiocyanate-dextran (FITC-dextran) was used to form maps of the healthy control angioarchitecture. C) Damaged cortical angioarchitecture following TBI at 25dpi corresponding to the same control region. D) Vascular topology at different cortical depths visualized at 25dpi after focal cortical injury. Left panel is a maximal projection image encompassing 250 μm of cortex that is color-coded for depths (top: red/pink; bottom: blue). The middle and right panel are images from the same brain injured mouse at different cortical depths, with more superficial cortical layers showing increased newly generated microvessels. E) Vessel density is quantified from 2P angioarchitectural maps, revealing varying vascular density with depth. At 23dpi there is an increase in vascular density at more superficial cortical layers (0-100 μm relative to the imaging start position) but appeared similar to pre-injury densities at deeper layers (101-200 μm relative to the imaging start position). The right panel shows averaged vessel density into 100 μm bins further documenting a significant increase at 23dpi compared to 19 and 39dpi but not pre-injury values (* p<0.05, repeated 2-way ANOVA). (Left panel: (n = 3) normalized to the first 20 μm cortical vessel density to facilitate comparisons across time).

Vessel density varies between pre-injury and at 19, 25, and 39dpi, with the highest pre-injury control vascular density at ~125μm into the cortex (Fig. 4E). Vessel density at 19 and 39dpi exhibit a similar depth profile. At 25dpi vessel density is up to ~50% greater relative to controls, particularly at upper cortical layers (Fig. 4E; see middle image Fig. 4D). Dichotomizing vessel density into < 100 μm and > 100 μm bins, confirms that at 25dpi there is a robust and significant increase in vessel density relative to 19 and 39dpi (Fig. 4E).

Overall results from *in vivo* 2P imaging of superficial cortical layers are consistent for most aspects of post-injury vascular remodeling compared to *in vivo* miniscope imaging (Figs. 2, 4D) and those from vessel painting (Figs. 1, 4D). One notable difference is that 2P imaging supports the notion that post-injury vascular pruning occurs at later post-injury time points (39dpi, Fig. 4E). In summary, our combined 2P imaging with miniscope imaging and vessel painting clearly demonstrates that the pre-injury vascular network is substantially altered following TBI. Small vascular loops are often observed as well as abnormal vessel plexuses with very fine vessels surrounded by larger blunted and incomplete vessels (Fig. 4D; Figs. 2E, 3A; Supplementary Fig. 1). Thus, while we clearly demonstrate vascular remodeling following TBI, the resulting angio-architectural restoration remains abnormal relative to control mice.

### Functional recovery of post-injury remodeled vascular networks

Angio-architectural remodeling of vessel networks implies that they may provide some level of physiological recovery of blood flow to injured tissues. To determine this, we assessed blood flow measures derived from miniscope experiments. We find significant loss of blood flow shortly after injury, which then undergoes progressive improvements in blood flow up to 33dpi, as shown by velocity maps (Fig. 5A). The qualitative observations are confirmed by longitudinal *in vivo* measurement of post-injury blood flow (Fig. 5B). Like vessel density, there is an initial peak in blood flow by 12dpi followed by a second peak in blood flow at 24-33dpi. Interestingly, while vessel area increases (Fig. 5C), there is no apparent relationship between peak blood flow speed and vessel area (Fig. 5D; R^2^=0.01).

**Fig. 5.**
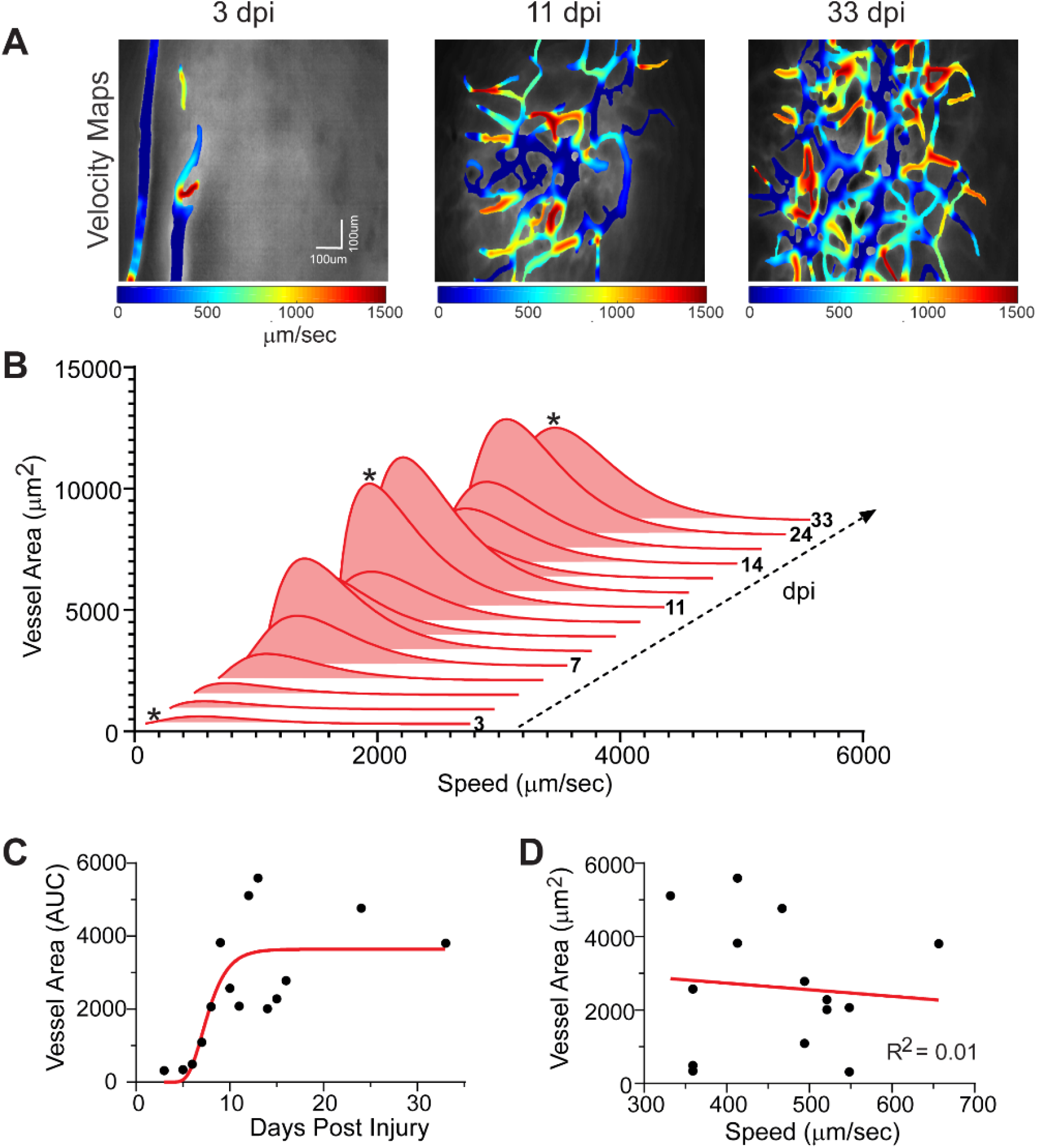
Temporal evolution of blood flow dynamics after the TBI injury. A) Color coded velocity maps at three representative time points after cortical injury (3, 11, 33dpi) further demonstrate increasing vascular density with attendant flow. B) Gamma distributions of blood flow velocity were obtained at each time point and the resultant histograms illustrate increasing flow velocity as the vascular networks increase in density. (* indicates the time points shown in A and C). C) Increased temporal vessel density at each time point (●) demonstrates vascular regrowth. Raw data is curve fitted (red line) using a Gompertz growth curve model. D) There is no correlation between maximal vessel area in the histogram and its associated blood flow velocity (R^2^=0.01). This suggests that as vessel density increases there is no overt change in the overall blood flow velocity. Similar findings were observed across all mice (n=4).

### Physiological regulation of blood flow returns

The head-mounted miniscopes enabled us to image blood flow changes within the same mouse under different behavioral states. We imaged vascular blood flow maps in TBI and control mice over the first two weeks mice. Representative flow maps obtained under three conditions, light (<1%) isoflurane anesthesia, awake and stationary, and running in their home cage are shown in Fig. 6A. At 6d post-TBI, the flow velocity is not regulated as the mouse went through the 3 activity states (Fig. 6B). In contrast, at 17dpi there is significantly improved vascular control in awake and stationary mice with decreased flow velocity compared with isoflurane anesthesia. The microvascular flow velocity increases when the mouse is running. This was further demonstrated by evaluating all control and TBI mice (Fig 6B, C). Under anesthesia there is no relationship to blood flow control and days post-injury (Fig. 6B; TBI R^2^=0.0832) but a stronger correlation is apparent while running with better vascular control at later time points (Fig 6C, R^2^=0.4154). Control mice exhibit no overt changes under any condition. There is a modest negative correlation between vessel diameter and flow velocity under anesthesia; larger diameter vessels show reduced flow (R^2^=0.524; Fig. 6D). Next, we binned TBI velocity data (n=3) into the closest control (n=4) measurement time point to examine the temporal relationship (Fig. 6E). Two-way repeated ANOVA reports a group effect (p<0.0001) but no significant temporal effect, whereas post-hoc testing reports significant differences at 4, 7 and 10dpi (p<0.0001). The longitudinal *in vivo* miniscope imaging demonstrates that the post-injury remodeled vascular network can respond physiologically, suggesting that some level of vascular control is present in the new vessel network. This finding could not be determined using static “snap-shot” measurements.

**Fig 6.**
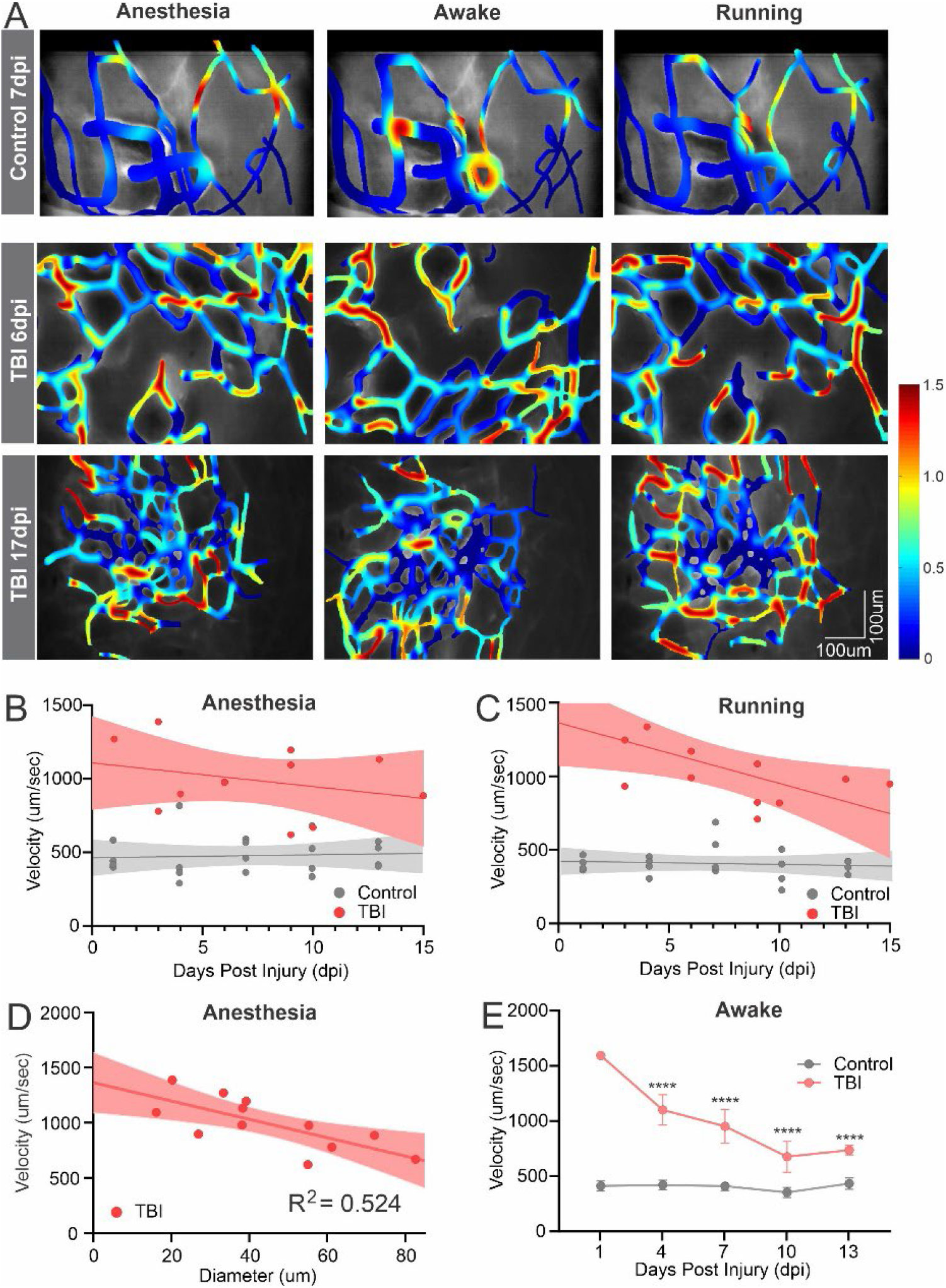
*In vivo* miniscope imaging reveals differential vascular regulation at multiple behavioral states at different time points following TBI injury. A) Exemplar color-coded plots of blood flow velocities from control and TBI mice (6and 17dpi) under isoflurane anesthesia, awake and running conditions. (Color scale codes the velocity in mm/sec). B) In TBI mice under anesthesia there was no relationship between microvascular blood velocity to days post injury (R^2^=0.0832). C) Under running conditions there was a stronger correlation between blood velocity and days after injury (R^2^=0.4154). D) Vessel diameter and blood flow velocity exhibited a modest relationship irrespective of days post injury (R^2^=0.524). E) There was a significant temporal decrease in blood velocity in TBI mice (n=3) that by 10dpi plateaued but never reached control mice (n=4) values (two-way repeated ANOVA group effect (p<0.0001) with significant differences (p<0.0001) at 4, 7 and 10dpi, post-hoc Tukeys) All linear regressions use the full data range to 15dpi and are plotted as mean and 95% CI.

### Endothelial cells guide new vessel sprouting to establish functional microvessels

Building on our findings of newly remodeled vascular networks, we probed the cellular processes underlying the initiation of new blood vessels and their functional engagement in blood flow. Endothelial cell (EC) guidance in vascular sprouting during angiogenesis plays a critical role in establishing functional perfused vascular networks following cortical injury. Specialized vascular ECs have distinct cellular fate specifications, including tip, stalk, and phalanx ECs, each having a unique role in vessel guidance and branching^25–28^. However, the early dynamic processes in response to adult cortical brain injury have not been studied *in vivo* due to lack of appropriate methodology.

The miniscope GRIN lens was implanted immediately following TBI (as in Fig. 2) and imaged daily over the first two weeks post-injury. This dynamic process is illustrated in an example mouse (n=3) (Fig. 7A-I). Starting at 3dpi, tdTomato-expressing ECs (red channel) are readily observed in the formation of new vessels (filopodia; Fig. 7C, red arrow) and they continue to grow progressively with increasing complexity throughout the measurement window of 2 weeks (Fig. 7A-I).

**Fig. 7.**
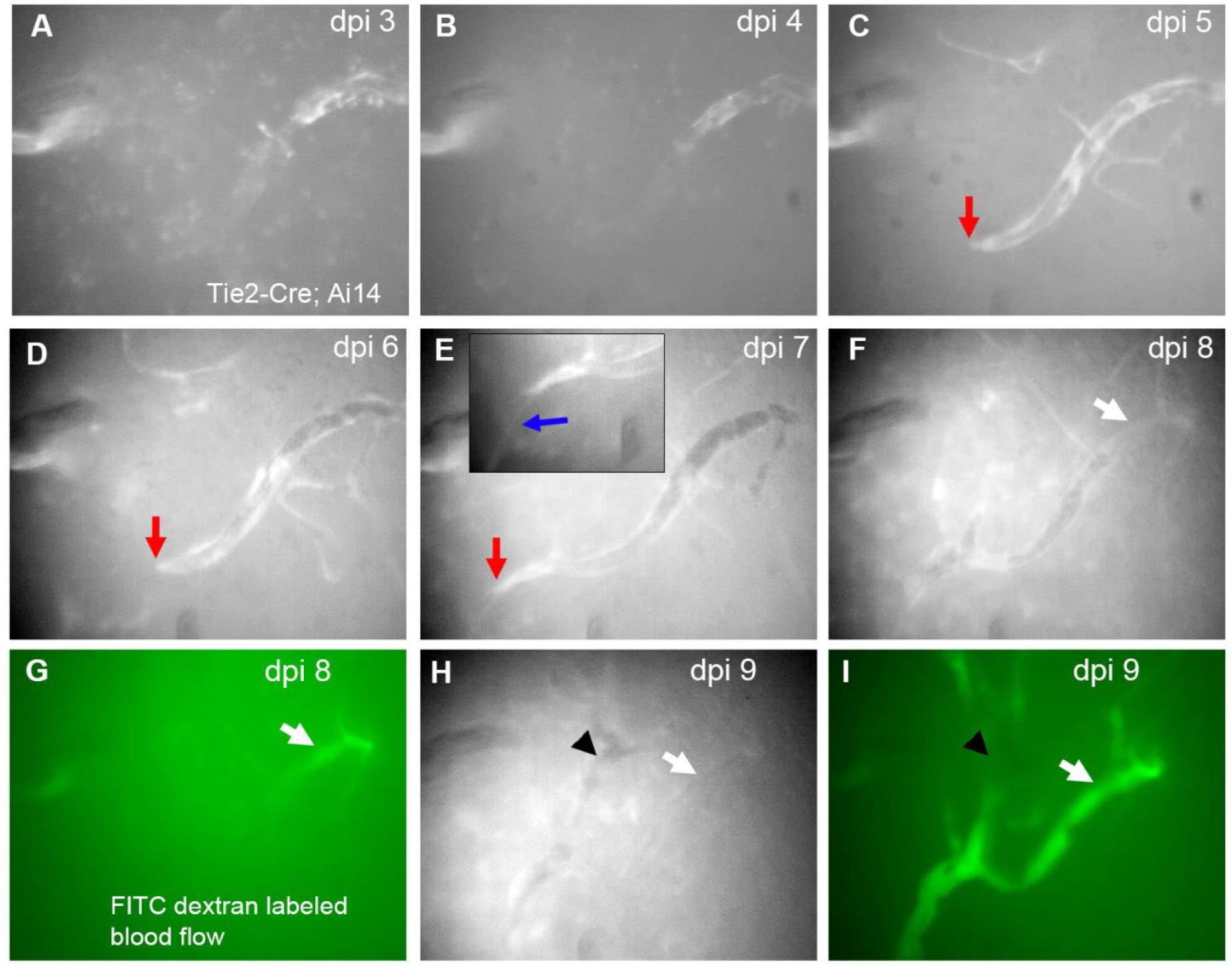
Dual-channel longitudinal *in vivo* miniscope imaging of endothelial cells and new vessel sprouting after cortical injury. The spatiotemporal pattern of ECs expressing tdTomato in a Tie2-Cre; Ai9 mouse was imaged daily at the same field of view in the injured cortex during the first 9dpi. Over the course of the first 8dpi (A-F), there is clear clustering of ECs in initial non-perfused microvessels, evidenced by intense fluorescent patchy regions in these vessels. Red arrows (C-E) point at a putative tip cell cluster that continues to migrate with increasing vessel length. The blue arrow (E) points to a filopodium emanating from the tip of the new micro-vessel. (G) and (I) illustrate a functionally perfused microvessel at 8dpi, visualized with FITC (green) fluorescent-labeled dextran for blood flow, which corresponds to the vessel (white arrows in F and G). (H) and (I) at 9dpi show increasing numbers of functionally perfused vessels. The arrows and arrowheads point to alignment features.

Dual channel miniscope imaging allowed assessment of functional perfusion of these tdTomato+ new blood vessels, with flows visualized by blood plasma labeled by FITC-conjugated dextrans (green channel) via tail vein injection at 8 and 9dpi (Fig. 7G, I; Supplementary Video 3). At 8dpi, only portions of the newly formed vasculature exhibit perfusion (Fig. 7G) but at 9dpi the same vessel and adjacent vessels are now being perfused (Fig. 7I) as the vascular network progressively remodels (Supplementary Video 3).

In a sub-cohort of mice, we performed 2P dual channel imaging to measure post-injury ECs, along with the fluorescein-labeled blood flow imaging (Fig. 8). In control mice, we detect individual ECs (red) that dot the surface of vessels (Fig. 8A, arrows). In the injury mice, at 4dpi nascent tube-like structures composed of ECs are visualized (Fig. 8B, arrowheads). However, these nascent vessels are not yet perfused as shown by the absence of fluorescein-labeled flow (green). In contrast, by 15dpi large vessels with abnormal aggregates of ECs are visualized at deep layers (~250um; Fig. 8C). Interestingly, at more superficial cortical layers, a dense vascular plexus of newly formed small vessels is very prominent (Fig. 8D), showing numerous EC-positive vessels but only a few of these small vessels support perfusion at this time point as shown by the relative absence of fluorescein-dextran signal.

**Fig. 8.**
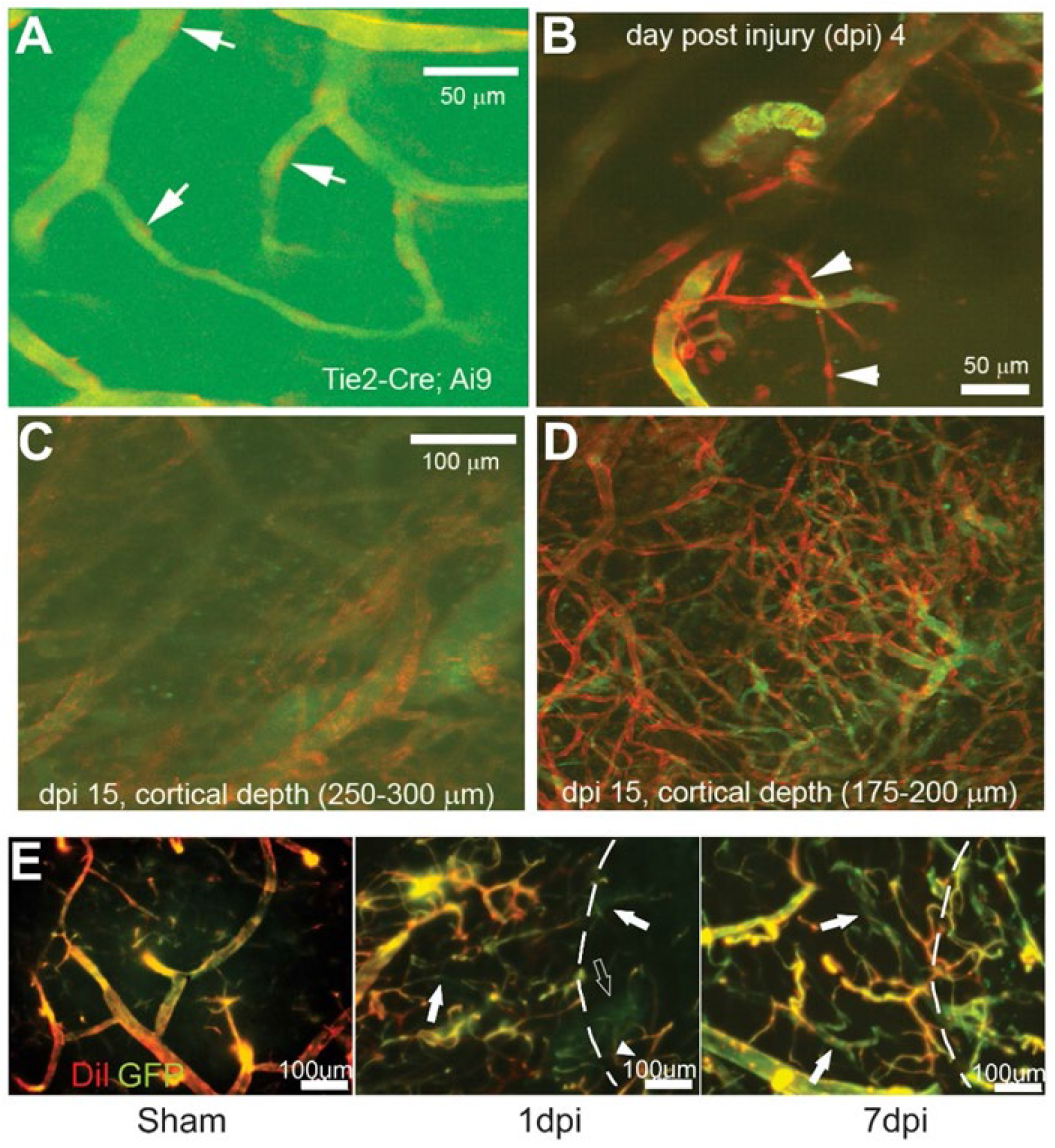
*In vivo* 2P imaging of new blood vessel sprouting and functional vascular networks after TBI injury. A) Visualization of tdTomato-expressing individual endothelial cells (arrows) in microvascular networks labeled by the fluorescein-dextran-stained blood plasma in a non-injured Tie2-Cre; Ai9 mouse brain. B) Consistent with *in vivo* miniscope imaging, 2P imaging identifies newly generated microvessels (tdTomato genetically labeled) in a Tie2-Cre; Ai9 mouse shortly (4 days) after injury. These vessels are not yet functional as they are not perfused with blood (FITC, green) (arrowheads). C-D) 2P imaging of the injury area at two cortical depths in a mouse at 15dpi. There were extensive newly re-generated microvessels at superficial cortical depths (175-200 μm; from cortical surface) that are not readily apparent at greater cortical depths. Note the tortuous nature of new microvessels. E) In sham control animals, Tie2-EGFP co-localizes with DiI (red) vessel painted vascular elements (left panel). At 1dpi there is a dramatic reduction in vessels in the injury zone (dotted line zone, middle) with new vessels (solid arrow) and newly perfused vessels (DiI+GFP, arrowhead) observable in the injury area. At 7dpi there is a dramatic increase in the number of perfused and non-perfused (arrows) vessels within and adjacent to the injury site (right). Note the increased density and tortuosity of the new vessels at 7dpi compared to sham.

The early temporal filopodial growth prompted us to further examine the progression of vascular growth from the early appearance of filopodia to the new vascular networks. Using *in vivo* 2P at 3d after TBI, a group of mice was sequentially imaged until 28dpi (Fig. 9). Like either vessel painting or miniscope imaging, early filopodia at 3dpi appear to underlie the development of local vascular networks (Fig. 9A). In several examples from different mice after TBI, we observe single filopodia that transform into dense vascular networks over the course of 7-10dpi (Fig. 9A-C). An important feature is that many of the new vessels that sprout from endothelial cell filopodia are poorly perfused or not perfused (Fig. 9A-C). At later time points after injury (~>14dpi) many of these emerging vascular elements are perfused; although in some TBI mice this functional recovery process takes longer (see Fig. 9C, 21dpi) with a dense vessel plexus (EC=red) that is not well perfused (FITC=green). Quantification of vessels that emerge from these filopodia shows that all mice exhibit a rapid increase in the vessel area (Fig.9D). In all mice, vessel density after TBI appears to follow a very similar pattern of rapid increases in density followed by a slower growth curve (Fig. 9E). These results provide further evidence that nascent vessel networks emerge from filopodia, then rapidly progress to increasingly functional vascular networks.

**Fig. 9.**
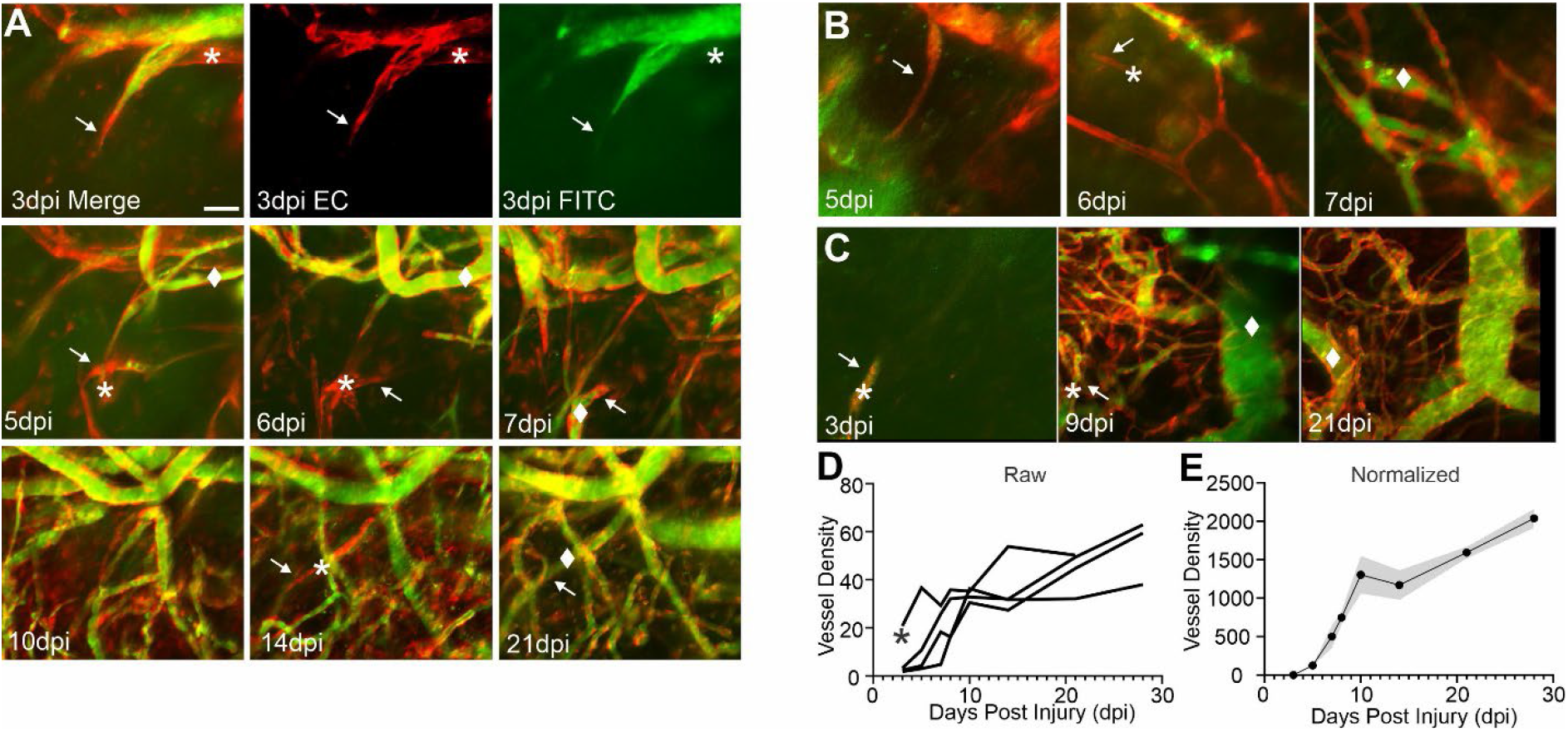
Emergent filopodia post cortical injury underlie re-growth of cortical vascular networks. A) Exemplar 2P images from the same mouse with longitudinal imaging over the course of 28d post TBI. Top panel shows the emergence of an endothelial cell filopodia (EC = red) (arrows) at 3dpi that is partially perfused (Green = FITC labeled blood flow). The * marks new vessel elements that are not yet perfused. As filopodia (arrows) continue to grow over the next several weeks, non-perfused vessels (*) increasingly become perfused as marked (◆). B) Another example from a mouse illustrating the rapid conversion of new filopodia (arrows) within 1 day from non-perfused (*) to perfused vessels (◆). C) In a third example, the left panel shows a single nascent filopodial element (arrow) at 3dpi that rapidly leads to a marked increase in vessel elements at 9dpi (center panel) that are predominately non-perfused (*). By 21dpi many of the vessels are more robustly delineated with endothelial cell bodies lining the vessels (right panel) and are being perfused (◆). D) Quantification of vessel density as a percent of the image area illustrates the progressive increase in vessel generation with time (n=4 mice). E) All raw image data are normalized to the first 2P recording (3dpi) to demonstrate the rapid growth of new vessels from nascent filopodia over the 28dpi recording period (mean ± SEM; n=4).

Similarly, in vessel painted brain tissues of Tie2-EGFP sham mice, GFP positive EC lumens co-localize with DiI-labeled vessels (Fig. 8E). At 1dpi there is a clear loss of vascular elements within the injury site as well as in the tissues adjacent to the injury. Some non-perfused vessels are already visible (Fig. 8E, 1dpi, white arrows) and the abnormal newly formed plexus of EC that are not perfused becomes visible (Fig. 8E, 1dpi, open arrows). By 7dpi, a robust increase in the number of vessels can be detected in the lesion although many remain un-perfused (Fig. 8E, 7dpi, white arrows). Therefore, as EC filopodia migrate to form new vascular networks, perfusion follows within the expanding network, thus potentially facilitating functional post-injury recovery of the surrounding brain tissues.

### Behavioral recovery coincides with vascular restoration

Behavioral and functional recovery following brain injury is an important consequence of vascular restoration. As our vascular imaging were mostly assessed after injury in the sensorimotor cortex, we conducted a series of behavioral experiments, primarily testing motor function, to determine if behavioral recovery coincides temporally with vascular restoration. Balance beam tests show progressive recovery with decreasing number of slips form 3 to 28dpi (Supplementary Fig. 2A). General exploratory activity shows that cortical brain injured mice spend significantly more time in the center of the arena at 14dpi, then return to control levels by 30dpi (Supplementary Fig. 2B). Foot-faults in post-injury mice while traversing a wire mesh peak at 3dpi, then rapidly recover coinciding with the time course of vascular remodeling (see Figs. 1, 2) (Supplementary Fig. 2C). The ability of mice to stay on a steady rotating cylinder is also impaired after injury, peaking at 3dpi but then rapidly recovers and functional recovery is sustained out to 45dpi (Supplementary Fig. 2D). While early sensorimotor deficits in mice after cortical injury are observed, most behavioral indices eventually return to control values, consistent with functional improvement in a similar time course as compared to vessel remodeling.

## Discussion

In this study, we applied new imaging technologies that we have developed to investigate CNS blood vessel formation and re-growth in response to a focal cortical injury. We report these novel findings. 1) There is a dramatic loss of post-injury cortical vessel density, which then progressively increases over the first 14-21d after injury. 2) This very dynamic vascular restoration can be visualized *in vivo* using miniscope and 2P imaging up to 60dpi. 3) Microvascular blood flow increases with increasing vessel restoration. 4) Vessel density within the cortex is most prominent in the more superficial layers, where thin and disorganized vessels are apparent. 5) Functional behavior recovers as vascular restoration proceeds. 6) *In vivo* imaging reveals new vessel sprouting within the injury site that is progressive and followed by attendant blood flow as EC filopodia expand. Our findings demonstrate the very dynamic and temporal nature of vascular restoration after cortical brain injury but also reveals that the resultant restored vessel network is structurally and functionally different from control animals. We show that microvessel remodeling occurs in cortical aspiration or contusion injury models. Broadly, our findings suggest a series of epochs, starting with a progressive increase in vessel density that peaks between 14-21d. This vessel remodeling results in vessel density like control and sham mice which is then followed by a wider distribution of blood flow within the nascent reconstructed vessel network. We and others have previously reported subacute (7-21d post injury) increases in vascular density in a variety of models of TBI^3, 5, 31^. Each of these studies reported significant increases in vascular density at 7-14d post injury relative to early injury time points. Park and colleagues in a fluid percussion brain injury model found no significant changes in cortical vessels at 14d post injury relative to shams, but the resultant vasculature was starkly abnormal5. Salehi et al3 at 7dpi and Haywood et al31 at 14dpi found increased vessel densities after cortical trauma, similar to our temporal findings. In focal photoembolic lesions of the cortex an identical growth of new vessels on the same time scale has been reported32, suggesting that similar mechanisms may underlie the brains ability to restore the vasculature.

Vascular pruning (regression) is a mechanism that has been described following proliferative angiogenesis in development^33, 34^, wherein the initial exuberance of vascular proliferation is now pared to a functional vascular density, as recently reviewed^34^. We observed a microvascular pruning process in histologically vessel painted brains and in *in vivo* miniscope and 2P microscopy imaging. The pruning of vessels appears to occur within 2-4 weeks after vessel regrowth in the two models we tested. Vascular pruning also occurs during postnatal development^33^, in aging^35^ and in other models of brain injury^32^. One functional rationale for pruning of excess vascular segments is that it serves to remove redundant and inefficient short vascular loops^36^. Indeed, in our miniscope imaging data we observed a proliferation of small vascular loops that were later pared to larger vessel loops. The precise signaling mechanisms for how these pruning events are achieved remain unknown; however now demonstrated in our study, these processes can be monitored in real time. Another phenomenon associated with pruning is the removal of vascular segments that become obstructed, where ~30% of obstructed vessels are removed^35^. Thus, dynamic pruning of the early vascular regrowth observed after brain injury serves to lead to a stable vessel network that can lead to a return of normal brain function even after brain injury.

Further evidence for dynamic microvascular remodeling was visualized using implanted miniscopes. We documented the formation of nascent EC filopodia at the injury site that progressively migrate into the injury site. The emergence of EC filopodia has been observed during development and is integral for formation of vessel networks^25, 37^. Several studies have shown *in explant* and *in vivo* studies of the retina, similar filopodia-driven revascularization after injury^38, 39^ and in the cerebral cortex in response to ischemic injury and tumor growth^40, 41^. We have previously documented filopodia and EC tip cell during re-vascularization after early after TBI^3^. Using Tie2-Cre; Ai9 mice we further demonstrated the emergence of robust new vessels at 4dpi that at superficial layers resulted in a large dense plexus of new vessels that were not yet perfused (Figs. 8, 9). Our findings show that filopodia directly lead to non-perfused vessel structures that become eventually become perfused as shown by FITC imaging. One notable feature is the appearance of a dense vascular plexus that remains without apparent flow even weeks after the initial TBI, either waiting to be perfused or is in the process of pruning. While not directly studied herein, in stroke these EC filopodia have been shown to extend to reactive astrocytes^42^. The importance of EC filopodia in vascular patterning has also been previously described^43, 44^. Finally, *in vivo* methods, such as miniscope and 2P imaging that allow repeated imaging of the vascular restoration process are highly amenable for tracking new filopodia growth and subsequent functional vascular networks to support “normalizing” blood flow.

The marked changes in vessel restoration we observe after cortical brain injury are accompanied by behavioral improvements. In a variety of behavioral paradigms, we find improved balance beam and open field performance over the course of 30dpi. The early recovery in foot-faults (3dpi) is potentially due to abatement of the initial insult of tissue edema, which follows a similar time course^45^. Many studies of cortical brain injury, such as stroke and TBI, have reported early recovery of deficits and been extensively reviewed^46, 47^. In a model of stroke, behavioral recovery corresponded strongly with blood flow adjacent to the injury site and related to vessel density^32^. In brain injury, others have reported an initial decrement in cerebral blood flow followed by an increase and then a later hypoperfusion epoch^31^. The temporal evolution decrease-increase-decrease is somewhat reminiscent of our vascular density findings that follow a similar pattern. Irrespective of the injury model, vascular restoration results in increased blood flow within the impacted tissues. We acknowledge that blood vessels are a single component of the neuron-astrocyte-blood vessel neurovascular unit and thus may be a contributory factor that together with enhanced neuronal and astrocyte health lead to behavioral improvements.

Together our study provides compelling novel data that following cortical brain injury, there is a very dynamic and robust growth of vessels to assist in functional tissue recovery. While there is significant functional (blood flow and behavior) post-injury recovery, it should be noted that many of the newly formed vessels at later time points (>30dpi) exhibit abnormal morphological characteristics that may compromise physiological function(s). There are a wide range of molecular and cellular mechanisms responsible for the brains attempt in repair. Developmentally, Wingless (Wnt) signaling is responsible for early tip cell and filopodia extensions in the brain^48^. Moreover, Wnt has also been implicated in healthy vascular remodeling^49^. Wnt mediates exuberant vessel growth in tumors^50^ and blockade of Wnt blunts tumor angiogenesis^51^. Stroke and brain injury studies have also focused on the role of vascular endothelial growth factor (VEGF) ^52–55^. We have previously suggested that Wnt is critical for initiation of vasculogenesis and VEGF (as well as other associated factors) is important for vascular stabilization^3^. Both Wnt and VEGF have a multitude of effects on the brain after injury including neurogenesis, gliogenesis and neuroprotection^56–58^. Extending the present study, future studies investigating how these molecular factors may play a role in the temporally dynamic restoration of the vessel network after cortical brain injury will improve the basis for the understanding of CNS microvessel repairs and for the treatment of cortical injuries.

## Abbreviations

2P: Two-photon
AP: Anterior-Posterior
CCI: Cortical contusion injury
CI: Confidence interval
dpi: days post injury
DV: Dorsal-Ventral
EC: Endothelial cell
FITC: Fluorescein isothiocyanate
GRIN: Gradient-index lens
ML: Medial-Lateral
PBS: Phosphate buffered saline
PFA: Paraformaldehyde
RBC: Red blood cell
SEM: standard error of the mean
TBI: Traumatic brain injury
VEGF: Vascular endothelial growth factor
Wnt: Wingless-Int1

## Acknowledgements

None

## Supplementary Materials

Supplementary materials include 3 supplementary videos and 2 supplementary figures.

Supplementary Video 1: Longitudinal miniscope imaging of microvascular recovery following cortical contusion injury.

Supplementary Video 2: Longitudinal miniscope imaging of microvascular recovery following aspiration cortical injury.

Supplementary Video 3: Dual-channel longitudinal miniscope imaging of newly generated microvessels after cortical injury.

