## Supplementary material for "Longitudinal dynamics of microvascular recovery after acquired cortical injury": Supplimental Figures

**Online Supplement for:** Longitudinal dynamics of micro-vascular recovery after acquired cortical injury.

**Supplemental Figures**

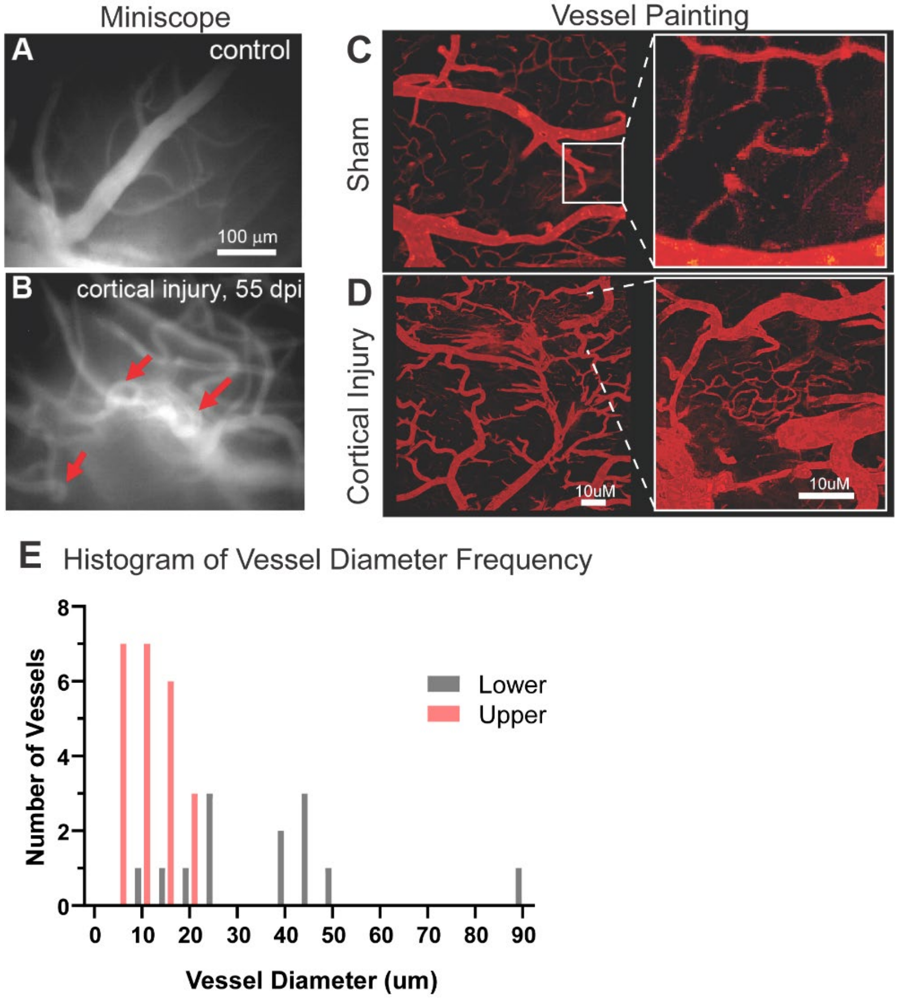


**Supplementary Fig. 1.** The restored microvascular networks exhibit abnormalities in vessel branching and organizational patterns. Miniscope Panel: A) A maximum intensity projection (MIP) image of the superficial cortical vessels in a control/sham animal (no injury, skull removed) with a GRIN lens at the cortical surface. The vessel branches remained stable for weeks. B) A MIP image of abnormal regenerated small blood vessels at 55dpi, where circular and loop-like structures formed (red arrows) in a TBI mouse. Vessel Painting Panel: C) Vessel painted cortical vessels in a sham operated mouse illustrate normal vascular structures, including normal capillary networks (see inset). D) Like GRIN lens imaging, at 30dpi the restored vascular network is abnormal and capillary networks are aberrant (see inset) after TBI. E) A frequency distribution histogram of vessel diameters derived from *in vivo* 2P imaging that illustrates that the upper cortical layers have a dense plexus of small diameter vessels whereas in the deeper cortical layers, larger diameter vessels are predominant (see Fig. 4).

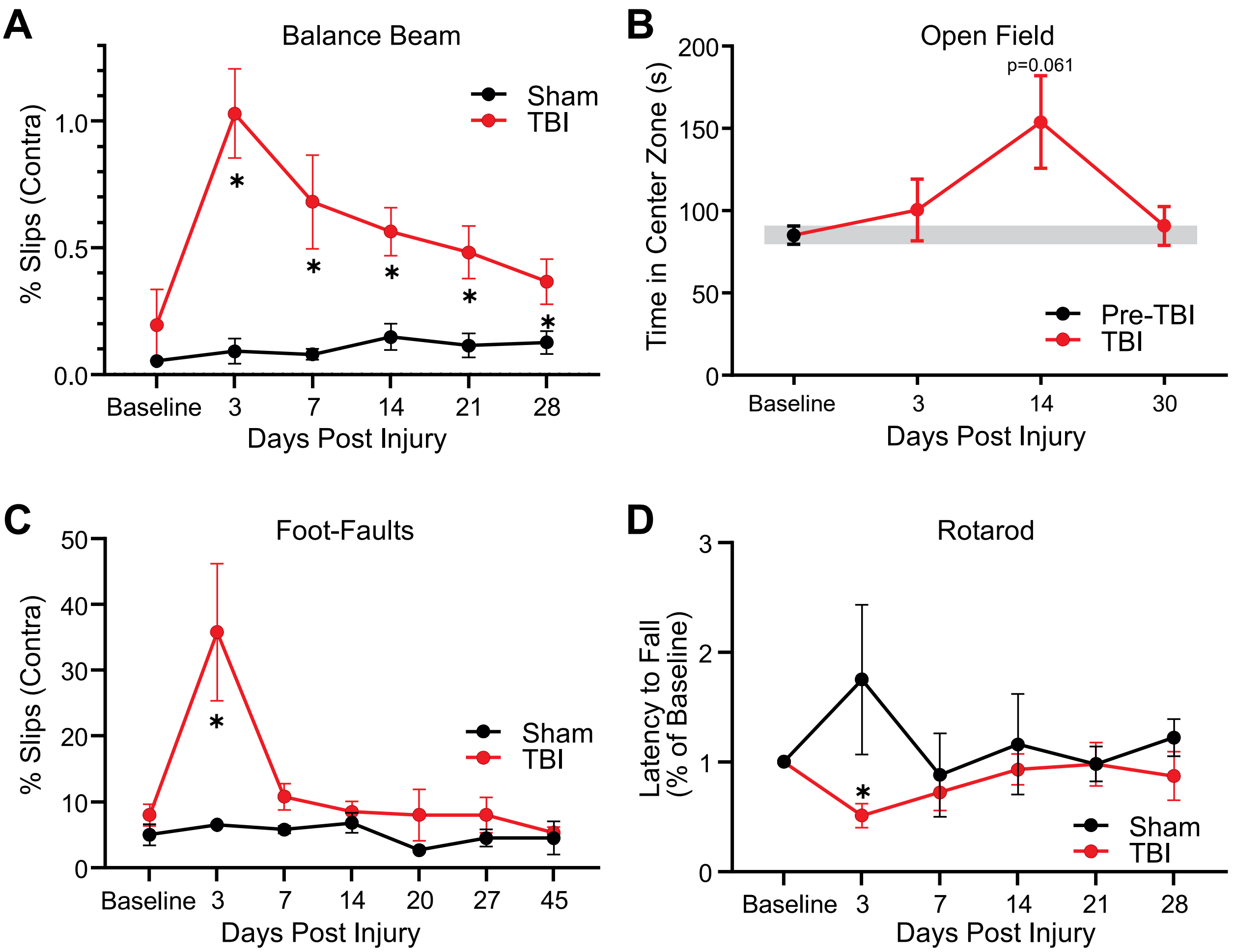


**Supplementary Fig 2. Behavioral assessments.** A) Sham (n=5) Tie2-Cre; Ai9 mice exhibited a low number of Beam Balance (BB) slips, while TBI mice (n=8) have a significant number of slips starting at 3-21dpi. B) Open field tests in cortical injury mice show a trend to spending increased time in the center zone at a time when vascular networks are most dense (14dpi) but return to baseline levels after vascular pruning is completed (n=4; One-way ANOVA with repeated measures). C) Foot faults (FF) on a wire mesh reports a significant increase on the side contralateral to the cortical injury (n=6; Two-way ANOVA with post hoc comparisons). D) Rotarod reports a decreased latency to fall at 3dpi that slowly recovered to baseline (n=3 shams, n=5 cortical injury; two-way ANOVA with post hoc comparisons) (* $p<0.05$)

**Legends for Video files**

Supplemental Video 1: Longitudinal miniscope imaging of microvascular recovery following cortical contusion injury (2, 6 and 22 dpi).

Supplemental Video 2: Longitudinal miniscope imaging of microvascular recovery following aspiration cortical injury (9, 12 and 62 dpi)

Supplemental Video 3: Dual-channel longitudinal miniscope imaging of newly generated micro-vessels after cortical injury (8 and 9 dpi).
